# Lense: Optimizing data preprocessing in single-cell omics using LLMs

**DOI:** 10.64898/2026.05.07.723465

**Authors:** Jingyun Liu, Zhicheng Ji

## Abstract

Data preprocessing is critical for single-cell omics analyses, but default pipelines often underperform on diverse datasets, especially from emerging platforms like spatial transcriptomics. We introduce Lense, a language-model-guided method that automatically selects optimal preprocessing by comparing plots that visualize low-dimensional representations across pipeline variants. Integrated with Seurat, Lense streamlines analysis and improves preprocessing robustness without requiring manual tuning.

**Biographical Note:** Jingyun Liu is a Master’s student in the Department of Biostatistics and Bioinformatics at Duke University. Dr. Zhicheng Ji is a tenure-track Assistant Professor in the Department of Biostatistics and Bioinformatics at Duke University. His research focuses on artificial intelligence and statistical modeling for single-cell genomics, spatial genomics, and biomedical imaging.

## Introductions

Single-cell RNA sequencing (scRNA-seq) and single-cell-resolution spatial transcriptomics (ST) platforms have become indispensable for dissecting cellular heterogeneity within complex tissues^1–5^. The reliability of downstream analyses often depends on a carefully engineered preprocessing pipeline that includes data normalization, selection of highly variable genes, dimensionality reduction, and cell clustering. Each stage offers multiple algorithmic alternatives and tunable parameters. For instance, data normalization can involve log transformation, centered-log-ratio scaling, or relative count rescaling. Although popular frameworks such as Seurat^6^ and Scanpy^7^ provide access to these options, analysts frequently adopt default settings, a practice that may lead to suboptimal performance.

The challenge is exacerbated when pipelines optimized for one modality, for example scRNA-seq, are applied without modification to data from emerging platforms such as the 10x Xenium ST system. Differences in data distribution between scRNA-seq and single-cell-resolution ST platforms^8^ increase the likelihood that parameter settings validated on hundreds of scRNA-seq datasets may perform poorly on ST data. In Figure 1a, we illustrate this discrepancy by simulating a 10x Xenium dataset from an annotated scRNA-seq sample (Methods), then evaluating the agreement between the resulting cell clusters and the inherited cell type annotations using the adjusted Rand index (ARI). The default scRNA-seq preprocessing parameters yield an ARI far lower than that obtained with an alternative parameter combination, suggesting that pipelines optimized for scRNA-seq do not generalize to ST data and that dataset-specific parameter tuning is essential for accurate cell clustering. However, identifying optimal parameter settings often involves a manual trial-and-error process, in which analysts test multiple combinations and visually compare the results of data processing and cell clustering. This approach is time-consuming and does not scale to large parameter spaces. Furthermore, the absence of a gold standard renders the process highly subjective. Analysts with limited experience may struggle to select appropriate parameters, leading to inconsistent and irreproducible results across research groups. Consequently, a systematic, data-driven strategy is required to pinpoint the optimal preprocessing pipeline rather than relying on intuition and ad hoc heuristics.

**Figure 1.**
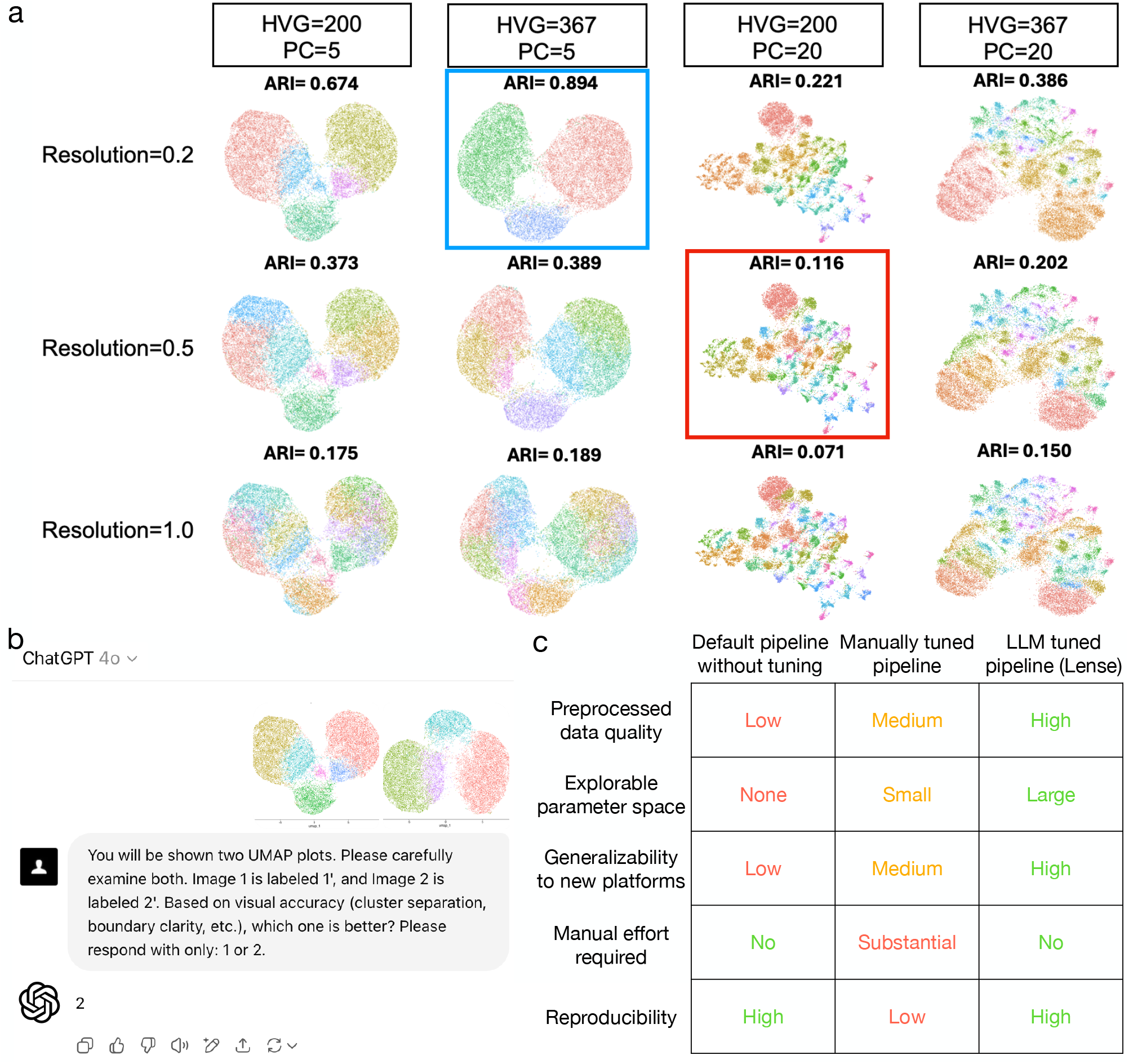
**a**, A simulated human pancreas 10x Xenium dataset demonstrating the effects of different preprocessing pipelines. The data normalization method used is log-normalization. Columns represent different numbers of highly variable features and principal components, while rows represent different cell clustering resolutions. ARI values are shown at the top of each subplot. The preprocessing pipeline closest to Seurat’s default setting is highlighted with a red rectangle, and the pipeline with the highest ARI is highlighted with a blue rectangle. **b**, A GPT-4o screenshot illustrating the prompt message and the output generated by GPT-4o for selecting the better UMAP plot. The ARI scores for the first and second UMAPs are 0.622 and 0.728, respectively. **c**, A table summarizing the advantages and limitations of existing methods and of Lense.

Large language models (LLMs), such as GPT-4 developed by OpenAI^9^, are advanced artificial intelligence systems trained on massive amounts of data and adaptable to a wide range of tasks. Although intended for general purposes, studies have shown that LLMs excel at analyzing genomic and biomedical data^10–14^. In this study, we present a strategy that uses LLMs to automatically select optimal preprocessing pipelines by evaluating plots that visualize low-dimensional representations, such as principal component analysis (PCA) or Uniform Manifold Approximation and Projection (UMAP), produced under different parameter settings. The rationale is that preprocessing quality is often visible in low-dimensional representations, where well processed data yields large, clearly separated clusters with few small, isolated islands. As illustrated in Figure 1b, GPT-4o was given two UMAPs generated from distinct pipelines and correctly identified the one with the higher adjusted Rand index (ARI), demonstrating its ability to recognize superior preprocessing configurations. Because generating low-dimensional representations is already a routine step of preprocessing pipelines and remains independent of the sequencing platform, this strategy avoids adding extra analysis steps, making it broadly applicable. By incorporating LLMs in place of human analysts, this strategy overcomes several critical limitations of existing methods (Figure 1c).

## Results

To systematically evaluate GPT-4o’s ability to identify the better preprocessing pipeline, we collected nine additional scRNA-seq samples generated and analyzed by different research groups and created simulated 10x Xenium datasets. Cell type annotations obtained from the original studies were treated as the gold standard. For each dataset, we applied three different cell filtering criteria with varying levels of stringency, resulting in a total of 30 scenarios. These scenarios represent diverse ranges in the number of cells, number of genes, average total number of reads per cell, and proportion of zeros (Figure 2a). We designed 72 preprocessing pipelines by combining six data normalization methods, two strategies for selecting highly variable features, two choices for the number of principal components, and three cell clustering resolutions. Each preprocessing pipeline was applied to every scenario, producing corresponding PCA and UMAP plots. A set of quantitative metrics, including ARI, Silhouette Score, Calinski–Harabasz Index (CHI), Davies–Bouldin Index (DBI), and Mutual Information (MI), was also calculated by comparing the cell clusters generated by each preprocessing pipeline with the ground truth. These metrics are used to determine which preprocessing pipeline yields results that better reflect the underlying biology and therefore represent a superior preprocessing strategy.

**Figure 2.**
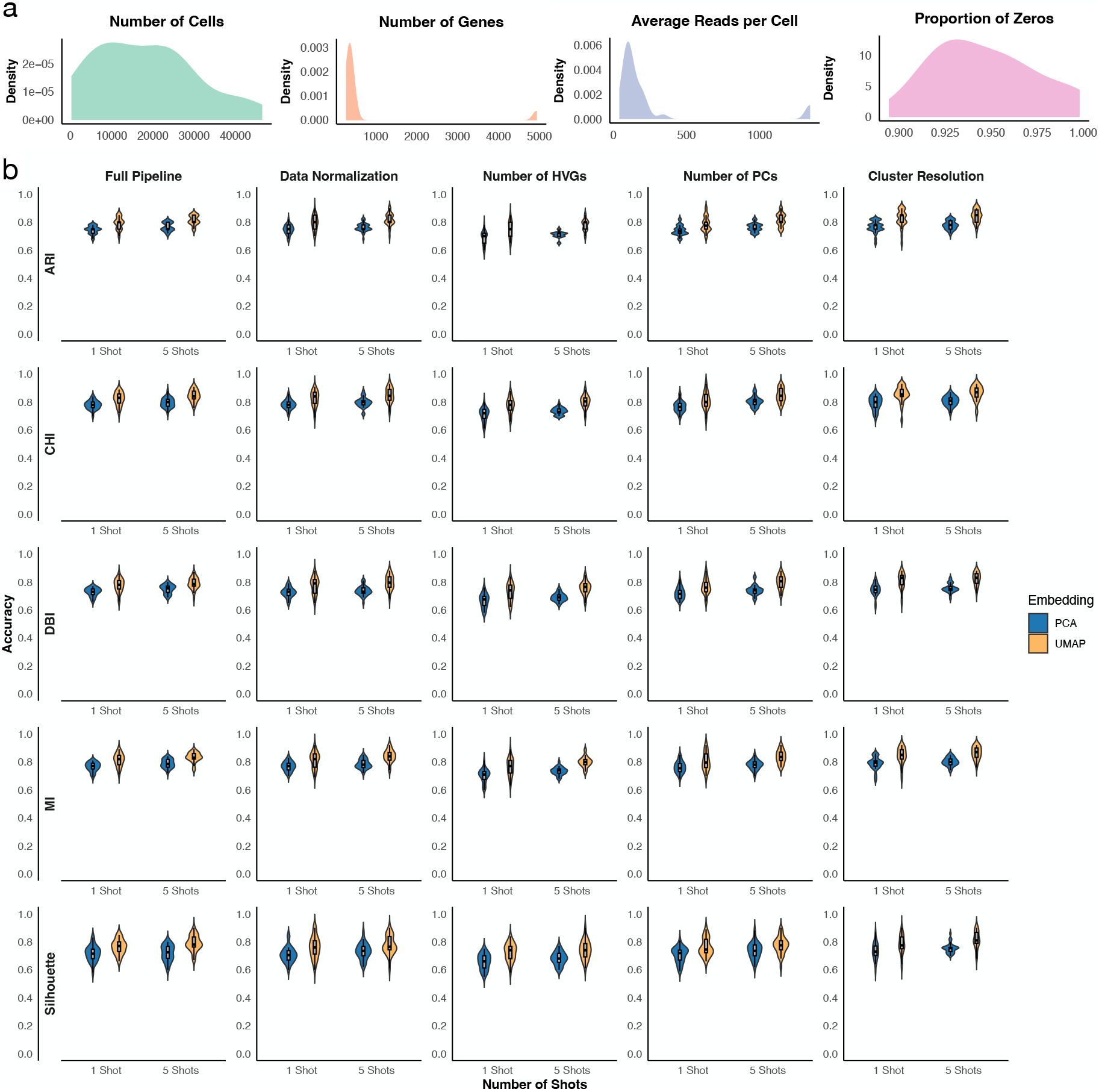
**a**, Distribution of the number of cells, number of genes, average total reads per cell, and proportion of zeros across the 30 simulation scenarios. Each data point represents one simulation scenario. **b**, Proportion of cases in which GPT-4o correctly identified the better pipeline according to a specific metric (row). Each data point represents one simulation scenario. The first column represents randomly chosen pairs, while the subsequent columns represent randomly chosen pairs that differ in only one preprocessing step, as indicated by the title.

For each scenario, we randomly selected 20 pairs of PCA or UMAP plots and prompted GPT-4o to choose the better one. Figure 2b shows that GPT-4o correctly identified the better preprocessing pipeline in most cases. For example, GPT-4o reached an accuracy of 80% using UMAP as input and ARI as the evaluation metric. The accuracy of GPT-4o increases slightly to 82% in a five-shot setting, where GPT-4o was prompted five times and the most frequent answer was used. GPT-4o remains highly accurate across evaluation metrics, and using UMAP as input shows slightly better performance than using PCA as input. We further performed a similar analysis on randomly selected pairs where only one of the four processing steps differed. GPT-4o demonstrated comparable one-shot and five-shot performance. These results suggest that GPT-4o can accurately and robustly identify the better preprocessing pipeline in pairwise comparisons. Note that UMAP and PCA plots are used solely as visual representations for pairwise comparison of preprocessing outcomes, rather than as quantitative validation metrics, and GPT-4o’s selections based on these embeddings show strong agreement with multiple quantitative metrics, indicating that they serve as informative proxies for relative comparison of preprocessing quality. Due to the limited improvement and increased computational burden of the five-shot approach, we used the one-shot approach in the subsequent analysis.

Based on these findings, we developed Lense, an LLM-powered preprocessing method for single-cell omics (Figure 3). Lense fully integrates with the Seurat pipeline and requires only one line of code to perform preprocessing. Lense first runs the Seurat preprocessing pipeline with different combinations of algorithms and parameter values. By default, 72 combinations described above are executed, but users have the option to include fewer or additional runs. A UMAP is generated for each run, and the corresponding preprocessed results are stored. Lense then iteratively prompts GPT-4o to perform pairwise comparisons of UMAPs and selects the best UMAP among the 72 candidates. Finally, Lense retrieves and returns the preprocessing results corresponding to the selected UMAP plot, which can be used for downstream analyses such as cell type annotation and differential analysis. Optionally, Lense can also generate PCA plots and select the optimal preprocessing pipeline based on these plots.

**Figure 3.**
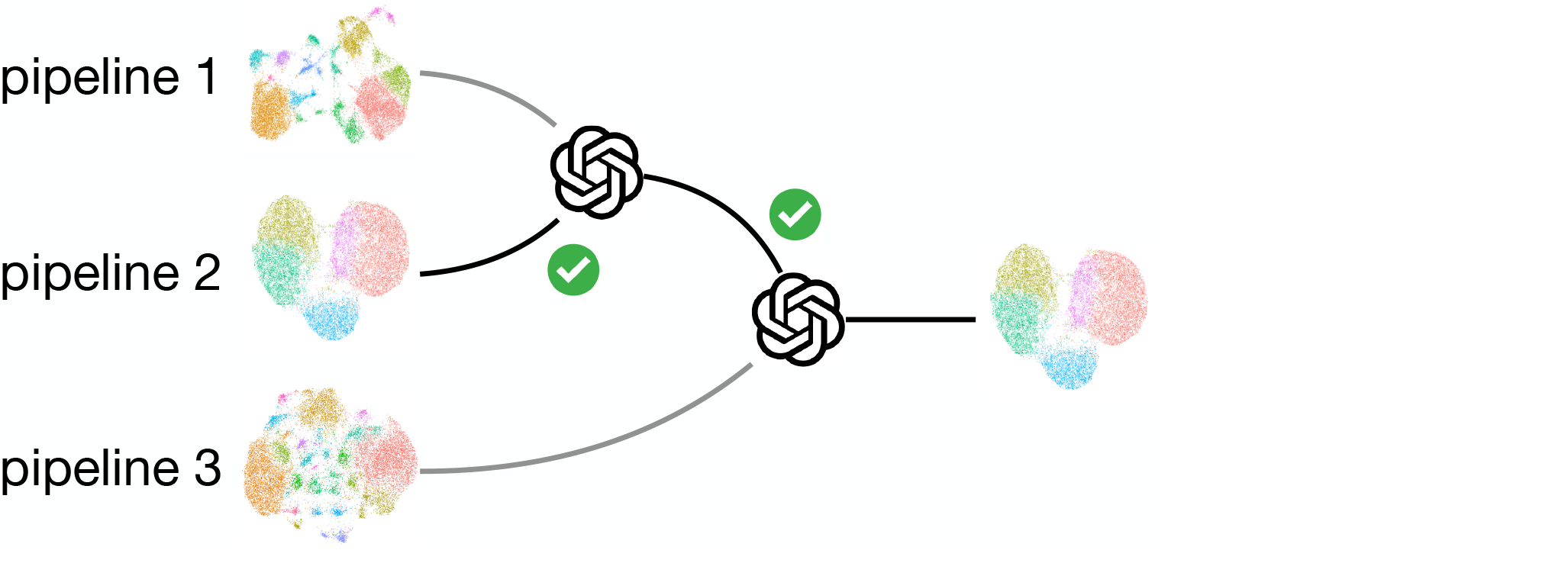
Cartoon illustrating the selection procedure of Lense.

We tested Lense on 30 scenarios and found that the preprocessing pipeline ranked as top by Lense consistently has among the best performance. For example, Lense with UMAP as input selected the preprocessing pipeline with the highest ARI in 73% of cases, and the selected preprocessing pipeline was among the top three in 90% of cases and among the top five in all cases (Figure 4). The results were similar when PCA was used as input to Lense or when other evaluation metrics were used. The overall performance of Lense is substantially better than that of any single fixed preprocessing pipeline and is very close to the theoretical optimum (“oracle”), where the preprocessing pipeline with the highest ARI or other evaluation metric was used for each dataset (Figure 5). These results suggest that a single fixed preprocessing pipeline is insufficient for handling datasets with highly diverse data distributions, and even the best-performing single pipeline may fail in certain cases (Supplementary Figure 1-2). In comparison, by adaptively selecting dataset-specific pipelines, Lense can effectively improve preprocessing efficacy.

**Figure 4.**
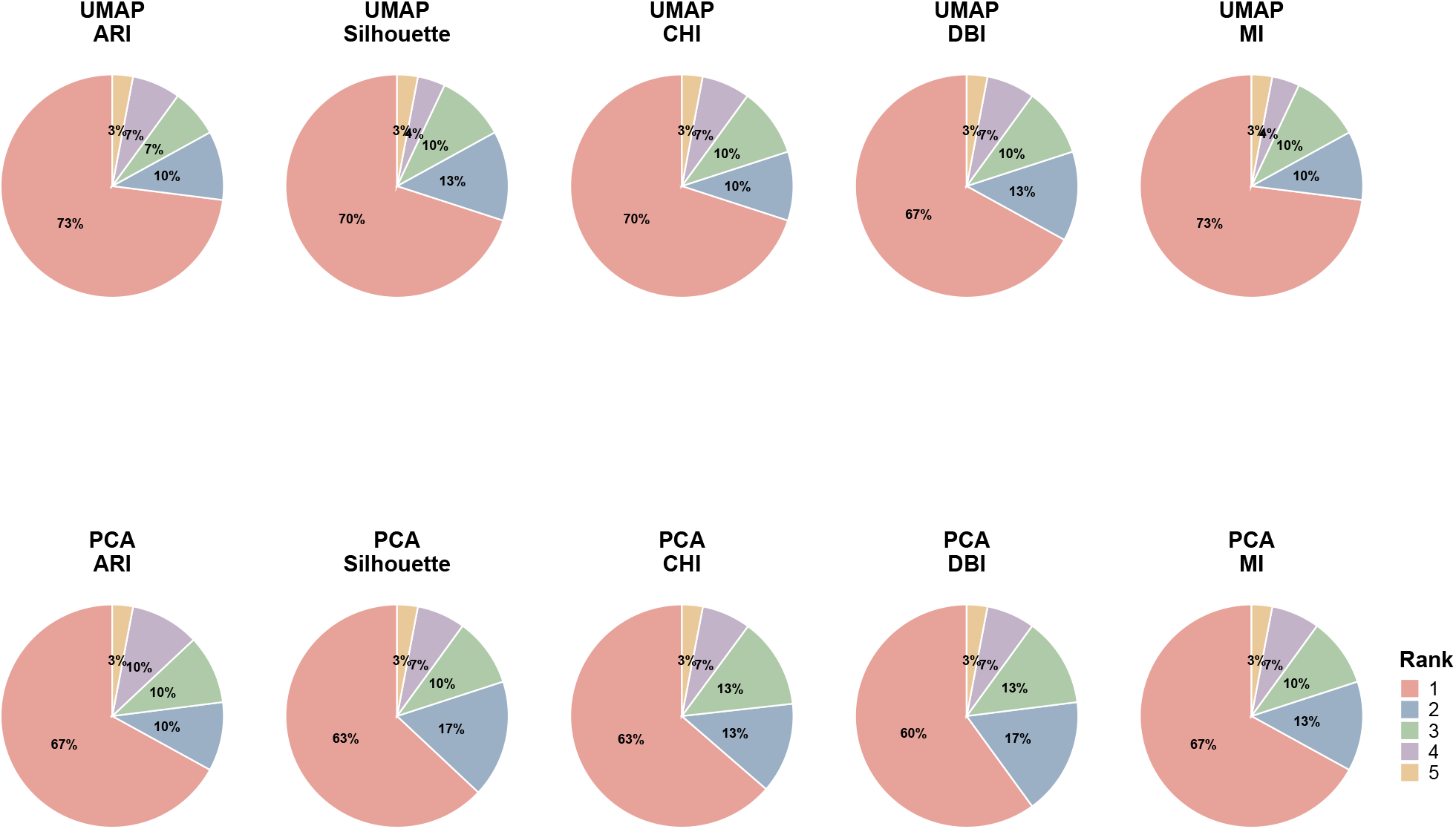
Proportion of simulation scenarios in which the pipeline selected by Lense achieved each rank according to a specific metric. The first row represents Lense using UMAP plots as input, and the second row represents Lense using PCA plots as input. Columns correspond to different evaluation metrics.

**Figure 5.**
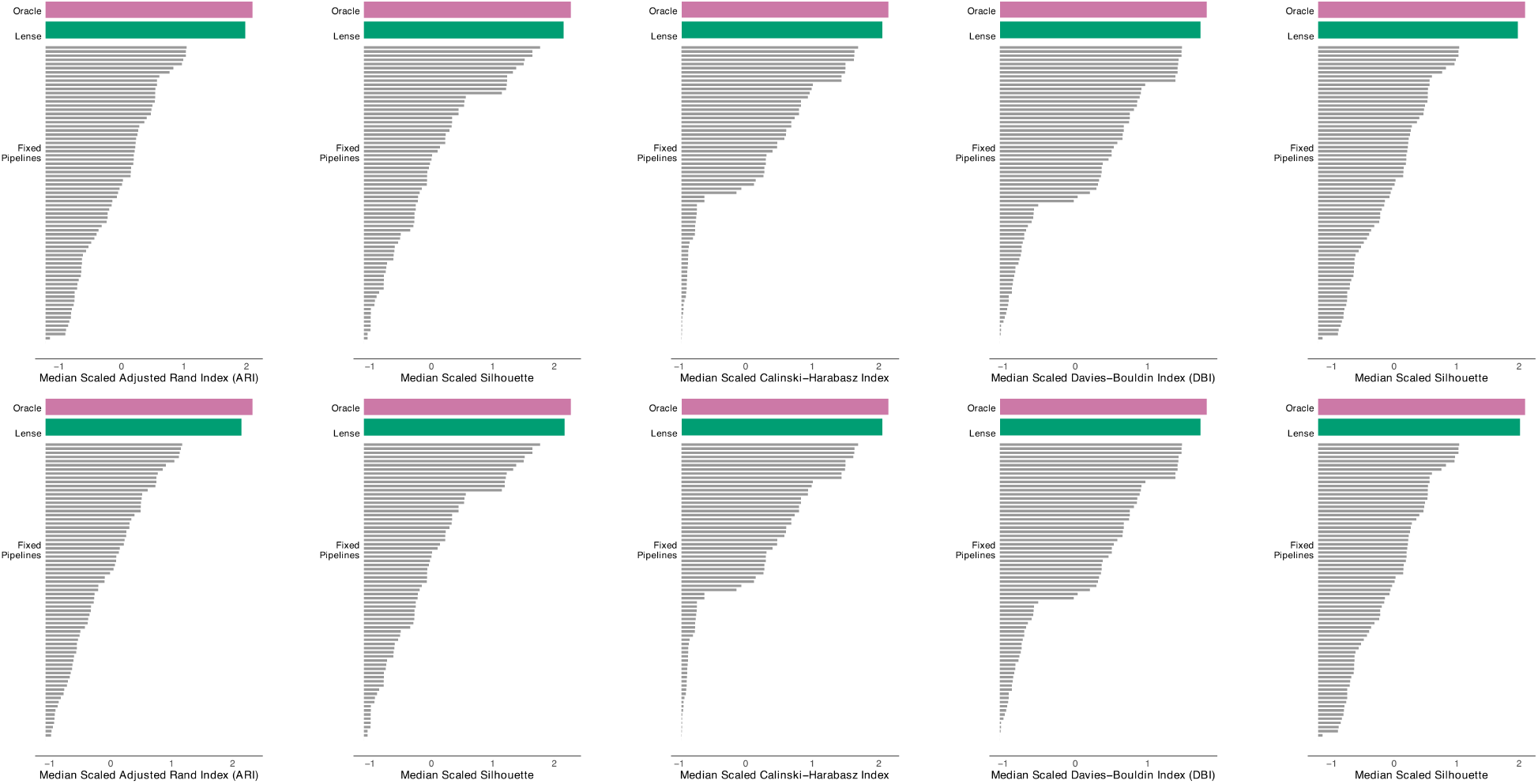
Medium scaled evaluation metrics for the theoretical optimum (“oracle”), Lense, and individual fixed preprocessing pipelines across scenarios. To make evaluation metrics comparable across scenarios, the original evaluation metrics were scaled to have a mean of 0 and a standard deviation of 1 across methods for each scenario. The first row represents Lense using UMAP plots as input, and the second row represents Lense using PCA plots as input. Columns correspond to different evaluation metrics.

We further evaluated the performance of Lense on two scRNA-seq datasets representing continuous differentiation processes (Figure 6). The first dataset, derived from human bone marrow, captures the differentiation of CD34+ hematopoietic stem cells. The second dataset, from the mouse dentate gyrus, represents neurogenesis. Using the same evaluation procedure, we compared Lense with single fixed pipelines across multiple evaluation metrics. The results show that Lense achieves the best performance in nearly all scenarios. The pipeline selected by Lense consistently ranks among the top three across both datasets, with either PCA or UMAP as input and across different evaluation metrics (Figure 6).

**Figure 6.**
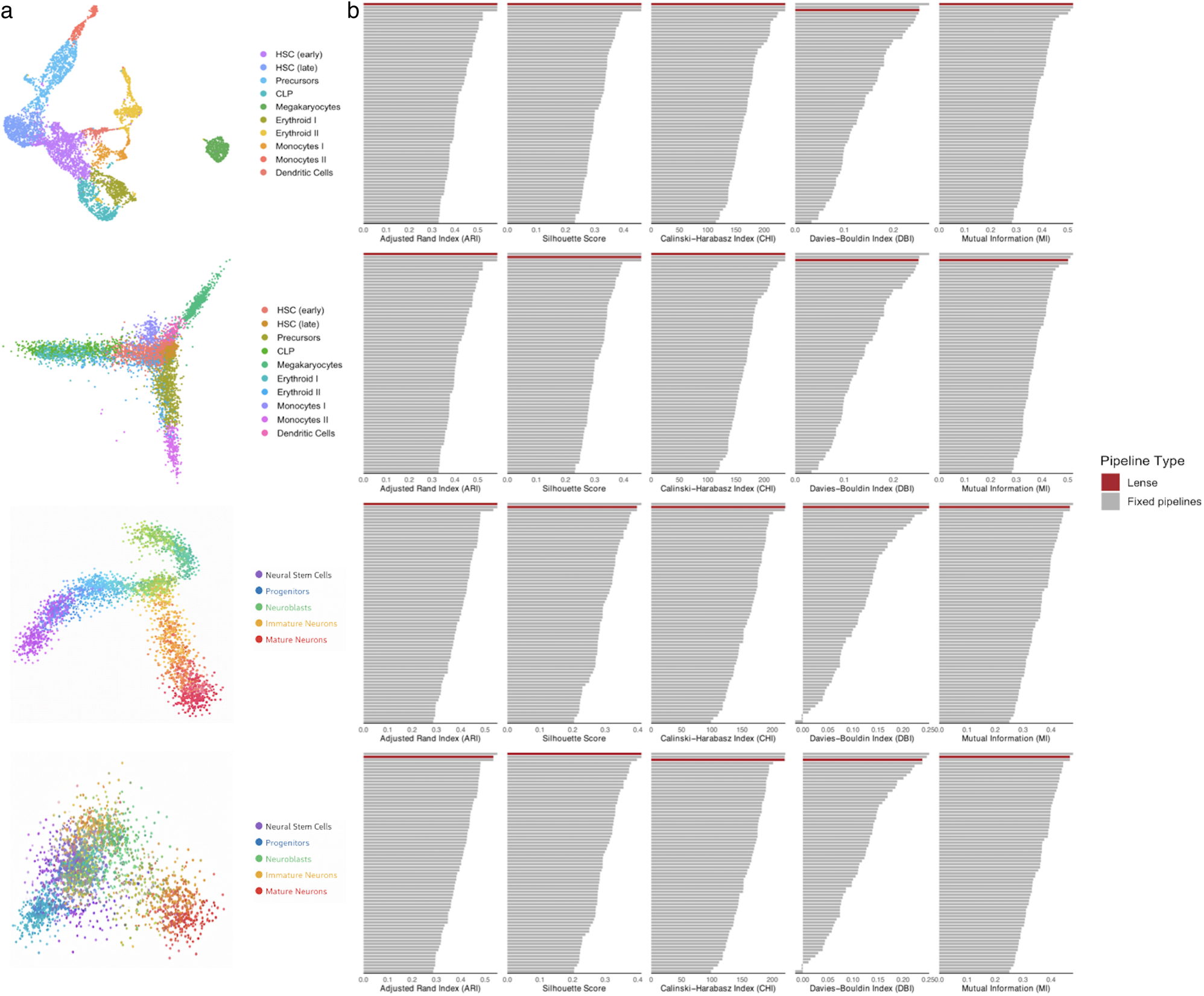
Performance of Lense on two scRNA-seq datasets representing continuous cell differentiation processes. From top to bottom: UMAP of the human CD34^+^ hematopoietic differentiation dataset, PCA of the same dataset, UMAP of the dentate gyrus neurogenesis dataset, and PCA of the dentate gyrus neurogenesis dataset. **a**, PCA or UMAP visualization of each dataset. **b**, Performance comparison between Lense and individual preprocessing pipelines.

Although LLMs can yield variable responses to identical prompts, Lense produces highly reproducible results. Across repeated runs, the Pearson correlation of pipeline rankings exceeds 0.9 in most cases (Supplementary Figure 3). We further assessed Lense using Claude and Gemini as alternative backbone LLMs instead of GPT-4o (Supplementary Figures 4–5). In these settings, Lense still ranks the optimal pipeline as the top choice in more than half of the cases and within the top five in nearly all cases. In addition, computational efficiency analysis shows that Lense scales linearly with the number of input cells and completes analysis within one hour for datasets of up to 20,000 cells (Supplementary Figure 6). Together, these results demonstrate that Lense is reproducible, robust, and scalable.

## Discussions

In summary, we developed Lense, which selects the optimal preprocessing pipeline by comparing PCA or UMAP plots generated from different pipelines. By formalizing preprocessing as a data-driven selection task enabled by high-level model reasoning, Lense provides a general framework for automating subjective steps in single-cell data analysis, with potential applications across diverse platforms and datasets.

The improved performance of Lense likely reflects not only its systematic exploration of a broad preprocessing space, but also its ability to harness the pattern-recognition capacity of large language models when applied to visual representations of data structure. Low-dimensional embeddings condense complex, high-dimensional transcriptional relationships into spatial configurations that encode cluster compactness, separation, continuity, and the presence of small, potentially artifactual islands. Human analysts routinely rely on such visual cues to assess preprocessing quality, yet these evaluations are subjective and difficult to standardize across datasets and research groups. In contrast, an LLM trained on large-scale visual and textual corpora may implicitly capture general principles of structural organization, enabling more consistent recognition of embeddings that display clearer inter-cluster separation, reduced fragmentation, and more coherent global geometry. In this way, Lense operationalizes visual intuition as an automated and reproducible decision process, allowing high-level visual reasoning to guide model selection across numerous candidate pipelines. The resulting performance gains suggest that LLM-based visual assessment can detect structural signals aligned with biological validity, underscoring a potential role for multimodal foundation models in the evaluation and optimization of analytical workflows beyond traditional quantitative metrics.

In this study, Lense was primarily evaluated using simulated 10x Xenium data, as the impact of preprocessing on ST data remains less well characterized than in scRNA-seq. Nevertheless, the data distributions in ST can differ substantially from those in scRNA-seq, and may also vary considerably across spatial platforms. Therefore, the extent to which these findings generalize to scRNA-seq or to other ST technologies warrants further investigation. In addition, although the simulation data were generated based on the real dataset, their distribution may still deviate from that of the actual data. This potential discrepancy should be considered when interpreting the results of Lense in real-world applications.

ST datasets often contain a large number of cells, and the original expression matrix is typically too large to be directly provided to LLMs. To efficiently convey the structure of the data, Lense leverages PCA or UMAP visualizations, which capture the low-dimensional embedding and clustering patterns of cells. Consequently, the performance of Lense may depend on the visual quality of the PCA or UMAP representation. For instance, visualization bias can arise when many points are plotted, as points rendered earlier may be obscured by those plotted later^15^. In addition, using similar colors for spatially adjacent clusters can make cluster boundaries difficult to distinguish^16^. These limitations may be mitigated by applying specialized visualization strategies^15,16^ to improve clarity and representation quality.

Lense is built upon the APIs of proprietary LLMs, such as GPT-4o, which currently demonstrate strong visual and reasoning performance relative to other available models^12^. However, this reliance on proprietary systems introduces limitations related to reproducibility, transparency, and long-term sustainability. These models are accessed through commercial APIs, while their underlying architectures, training data, and update schedules are not publicly disclosed. Consequently, model behavior may evolve over time without explicit version control, potentially affecting the stability and exact reproducibility of previously reported findings. In addition, the inference costs associated with commercial LLM usage can be substantial, particularly for large-scale analyses. Such financial and infrastructural demands may restrict accessibility for research groups with limited resources.

## Methods

### Lense

Lense consists of two steps. In the first step, it preprocesses the input single-cell omics data using the Seurat pipeline with various combinations of parameter settings. In the second step, it uses GPT-4o to compare the PCA or UMAP plots generated in the first step and selects the optimal preprocessing pipeline. By default, UMAP plots are used in the second step of the pipeline.

#### Step 1: preprocessing with different parameters

The input to Lense is a Seurat object that contains a count matrix and has undergone appropriate cell filtering. The data can originate from various single-cell sequencing platforms, such as scRNA-seq or spatial transcriptomics technologies with single-cell resolution. Lense then sequentially performs the following preprocessing steps using the Seurat pipeline: data normalization, identification of highly variable features, data scaling, principal component analysis, selection of the number of principal components, cell clustering, and visualization.

By default, Lense evaluates 72 distinct preprocessing pipelines derived from combinations of the parameters described below. Users may optionally select a subset of these pipelines or define additional ones.

##### Normalization methods

One of six normalization approaches is selected:

- SCTransform (SCT)^17^, implemented via the `SCTransform` function in the `sctransform` package
- Log-normalization (Lognorm): total read count normalization followed by log-transformation, implemented via `NormalizeData` in the `Seurat` package
- Centered log-ratio (CLR): implemented via `NormalizeData` in `Seurat`
- Relative counts (RC): total read count normalization, implemented via `NormalizeData` in `Seurat`
- Log transformation: log-transformation, implemented manually
- Raw counts: no normalization applied

##### Highly variable feature selection

One of two options is selected:

- Using all *I* features
- Selecting *a* highly variable features using the `FindVariableFeatures` function in `Seurat` with default settings, where

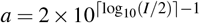

##### Number of principal components

The number of principal components (PCs) is set to either 5 or 20.

##### Clustering resolution

Cell clustering is performed using the `FindClusters` function in `Seurat`, with resolution parameters set to 0.2, 0.5, or 1.

All other steps follow the default parameters of the `Seurat` pipeline. Lense executes each preprocessing pipeline and records both the preprocessing results and the corresponding PCA or UMAP plot.

### Step 2: selecting the optimal preprocessing pipeline

Lense then performs pairwise comparisons to select the optimal preprocessing pipeline. Suppose there are *N* preprocessing pipelines generated in the first step. Lense conducts *N* − 1 pairwise comparisons sequentially. In the first comparison, Lense compares the first and second pipelines. In the *i*th comparison, Lense compares the (*i* + 1)th pipeline with the pipeline selected in the (*I* − 1)th comparison.

In each pairwise comparison, Lense uses the application programming interface (API) provided by OpenAI to query the gpt-4o-2024-08-06 model. Two low-dimensional embeddings, generated using either UMAP or PCA, along with the following text prompt, are used as inputs: “You will be shown two UMAP plots. Please carefully examine both. Image 1 is labeled ‘1’, and Image 2 is labeled ‘2’. Based on visual accuracy (cluster separation, boundary clarity, etc.), which one is better? Please respond with only: 1 or 2.” Due to the randomness and lack of controllability in the output of LLMs, the gpt-4o-2024-08-06 model is prompted repeatedly until its response is either “1” or “2”.

### Collection of matched spatial and single-cell omics datasets

We collected the gene expression count matrices of spatial transcriptomics samples generated using the 10x Genomics Xenium In Situ platform (https://www.10xgenomics.com/datasets/) from ten tissue types, including human kidney, human liver, human pancreas, human lung, human bone marrow, human brain, human breast, human lung cancer, human heart, and mouse brain.

For each tissue type, we also collected the gene expression count matrix and the manually annotated cell type labels from a publicly available scRNA-seq dataset generated from the same tissue. Specifically, human liver data^18^ were obtained from the Gene Expression Omnibus (GEO) under accession code GSE192742, human pancreas data^19^ from GEO under accession code GSE221156, human bone marrow data^20^ from GEO under accession code GSE166895, human lung cancer data^21^ from ArrayExpress under accession codes E-MTAB-14560 and E-MTAB-14566, adult human heart data^22^ from the European Nucleotide Archive (ENA) under accession code ERP123138, human breast data^23^ from ArrayExpress under accession code E-MTAB-13664, human lung data^24^ from GEO under accession code GSE161382, mouse brain data^25^ from CELLxGENE Discover platform^26^, human kidney data^27^ from GEO under accession code GSE202109, and human brain data^28^ from GEO under accession code GSE199762.

For scRNA-seq datasets that only provide Ensembl gene IDs, we used the biomaRt package^29^ to convert the gene IDs into gene symbols. Genes without valid gene symbols or with duplicated mappings were removed.

### Simulation of 10x Xenium datasets

A simulated 10x Xenium dataset was generated for each pair of real scRNA-seq and real 10x Xenium samples from the same tissue. Only genes shared between the two datasets were retained. A scaling factor was computed as the ratio of the mean library size of the real Xenium data to that of the scRNA-seq data. For each cell in the scRNA-seq dataset, a target library size was calculated by multiplying the cell’s total read count by the scaling factor. The reads in that scRNA-seq cell were then randomly subsampled so that the total read count matched the target library size.

### Collection of scRNA-seq datasets representing continuous differentiation processes

Two scRNA-seq datasets representing continuous differentiation processes were collected. The first dataset corresponds to human CD34^+^ hematopoietic differentiation. The raw gene expression count matrix was obtained from the Human Cell Atlas (HCA) data portal under project ID 091cf39b-01bc-42e5-9437-f419a66c8a45, which is associated with the study by^30^. This dataset comprises scRNA-seq profiles of hematopoietic stem and progenitor cells and captures continuous differentiation trajectories across multiple lineages. Cell type annotations were obtained from the original dataset metadata.

The second dataset corresponds to dentate gyrus neurogenesis and was obtained from the Gene Expression Omnibus (GEO) under accession code GSE95753^31^. This dataset contains scRNA-seq profiles from mouse dentate gyrus across multiple developmental stages, spanning embryonic to postnatal time points, and captures the progression of neurogenesis. Cell type annotations were obtained from the original study.

For both datasets, raw gene expression count matrices were converted into Seurat objects and used as input to the Lense pipeline, ensuring consistent preprocessing across all datasets.

### Evaluation of clustering performance

Clustering performance was evaluated by comparing the cell clustering results from a preprocessing pipeline with the ground truth cell type annotations inherited from the scRNA-seq dataset. Five quantitative metrics were computed: Adjusted Rand Index (ARI), Silhouette Score, Calinski–Harabasz Index (CHI), Davies–Bouldin Index (DBI), and Mutual Information (MI). ARI was computed using the `adjustedRandIndex` function from the mclust package in R. The Silhouette Score was calculated using the `silhouette` function from the `cluster` package, based on Euclidean distances in the reduced dimensional space. The Calinski–Harabasz Index (CHI) and Davies–Bouldin Index (DBI) were computed using the `cluster.stats` function from the `fpc` package. Mutual Information (MI) was calculated using the `mutinformation` function from the `infotheo` package. Since DBI is a metric where lower values indicate better clustering performance, we transformed it as 1− DBI so that higher values consistently correspond to better performance across all metrics.

### Additional LLMs

In addition to the default OpenAI GPT-4o model, we evaluated Lense using Claude 3.5 Sonnet (Anthropic) and `Gemini 1.5 Pro` (Google). All models were accessed via their respective APIs and applied under the same protocol to select optimal preprocessing pipelines based on visual comparisons of low-dimensional embeddings (UMAP or PCA).

## Key points

- Default preprocessing often underperforms on diverse datasets, especially single-cell-resolution spatial transcriptomics, so dataset-specific tuning is needed.
- Lense uses an LLM to compare PCA or UMAP plots from many pipeline variants, then selects the best preprocessing configuration automatically.
- Integrated with Seurat and callable in one line, Lense replaces manual trial-and-error, improving consistency and reproducibility without extra bespoke steps.
- GPT-4o correctly picked the higher-ARI UMAP in about 80% one-shot and 82% five-shot tests across 30 scenarios, showing robust visual-quality judgment.
- On the same scenarios, Lense chose the true best pipeline in 73% of cases, was top-3 in 90% and top-5 in 100%, performing far better than any fixed pipeline and close to an oracle.

## Acknowledgments

The project was supported by the National Institutes of Health under Award Number U54AG075936 and R35GM154865.

## Author contributions

Z.J. conceived the study. J.L. conducted the analysis and developed the software. Z.J. and J.L. wrote the manuscript.

## Competing interests

All authors declare no competing interests.

## Data availability

Lense is freely available on GitHub: https://github.com/jingyun20/Lense.

## Notes

### Competing Interest Statement

The authors have declared no competing interest.

