## Supplementary Figure for "Lense: Optimizing data preprocessing in single-cell omics using LLMs"

### Supplementary materials

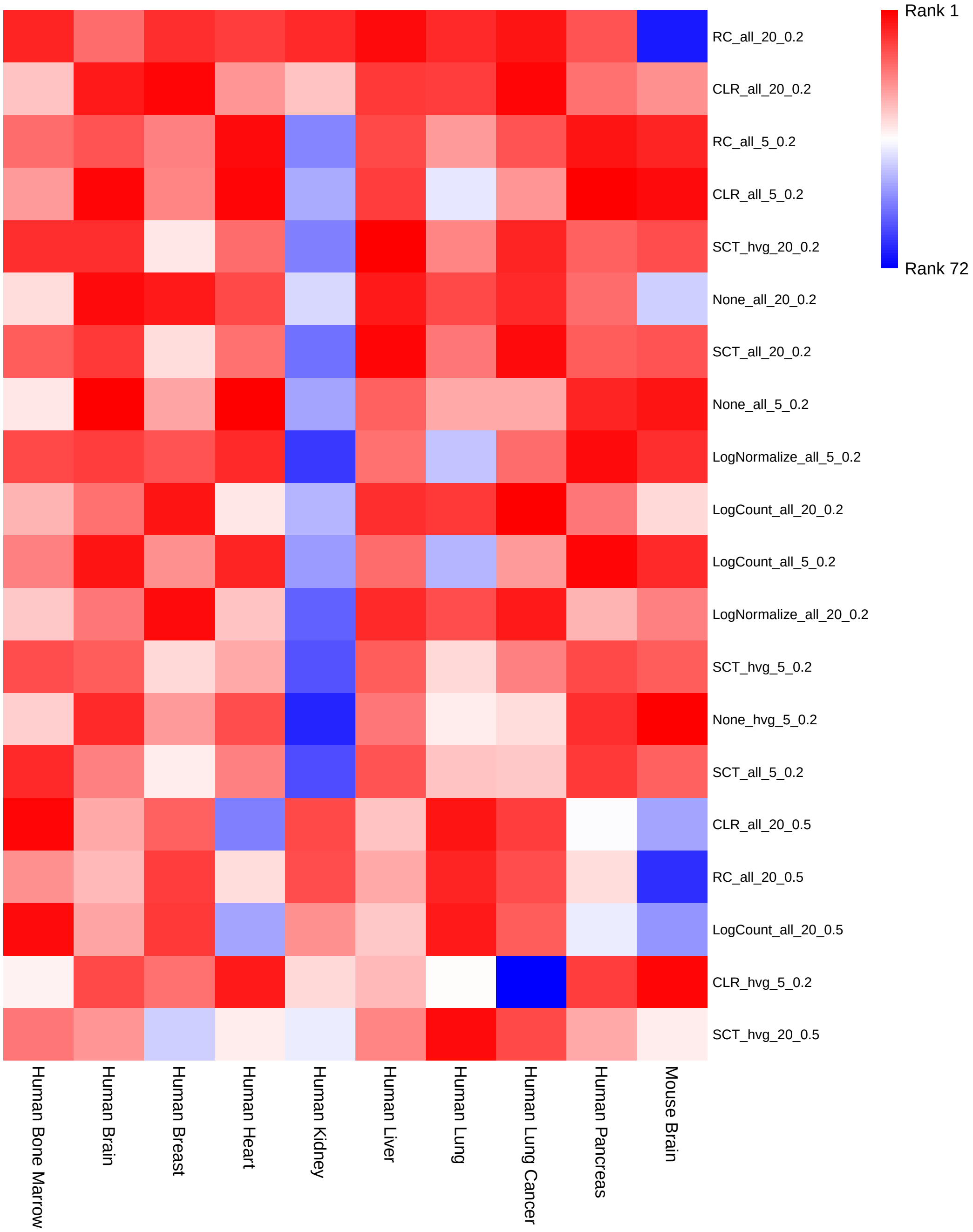


**Figure S1.** Ranking of top 20 best performing preprocessing pipelines in each dataset. Rows represent pipelines and are ordered based on their average rank based on ARI across datasets. Columns correspond to different datasets. Each cell shows the rank of a pipeline within a dataset based on ARI, where lower ranks indicate better clustering performance. Each pipeline is denoted in the format *Normalization_FeatureSelection_PC_Resolution*, where *Normalization* specifies the normalization method, *FeatureSelection* indicates whether all genes ("all") or highly variable genes ("hvg") are used, *PC* denotes the number of principal components, and *Resolution* represents the clustering resolution parameter. For example, *RC_all_20_0.2* represents a pipeline that applies relative count normalization, uses all genes, retains 20 principal components, and uses a clustering resolution of 0.2.

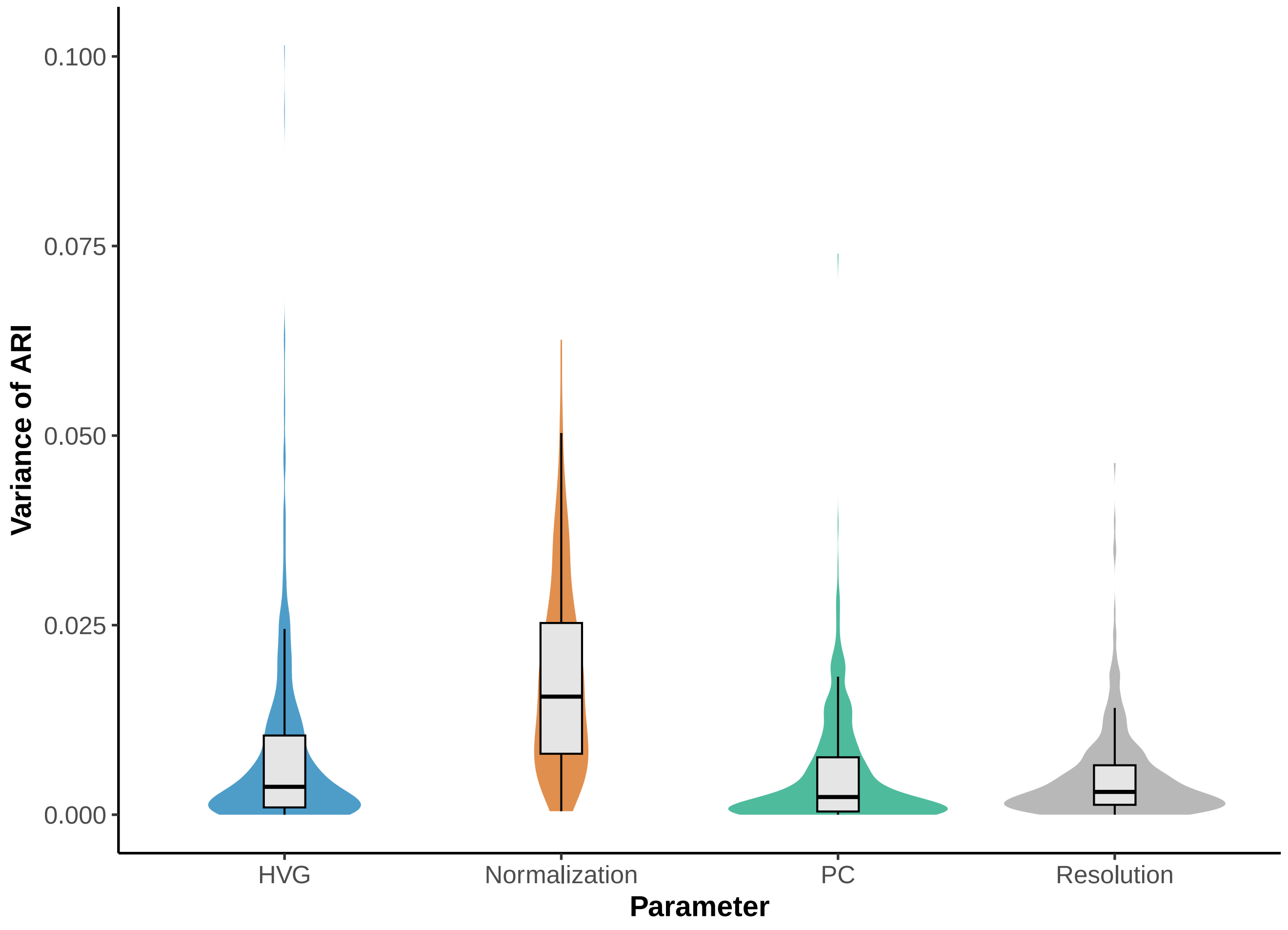


**Figure S2.** Variance in clustering performance across preprocessing parameters. Each violin plot represents the distribution of ARI variance for a given parameter setting, computed by fixing that setting and evaluating variance across all possible combinations of the remaining parameters.

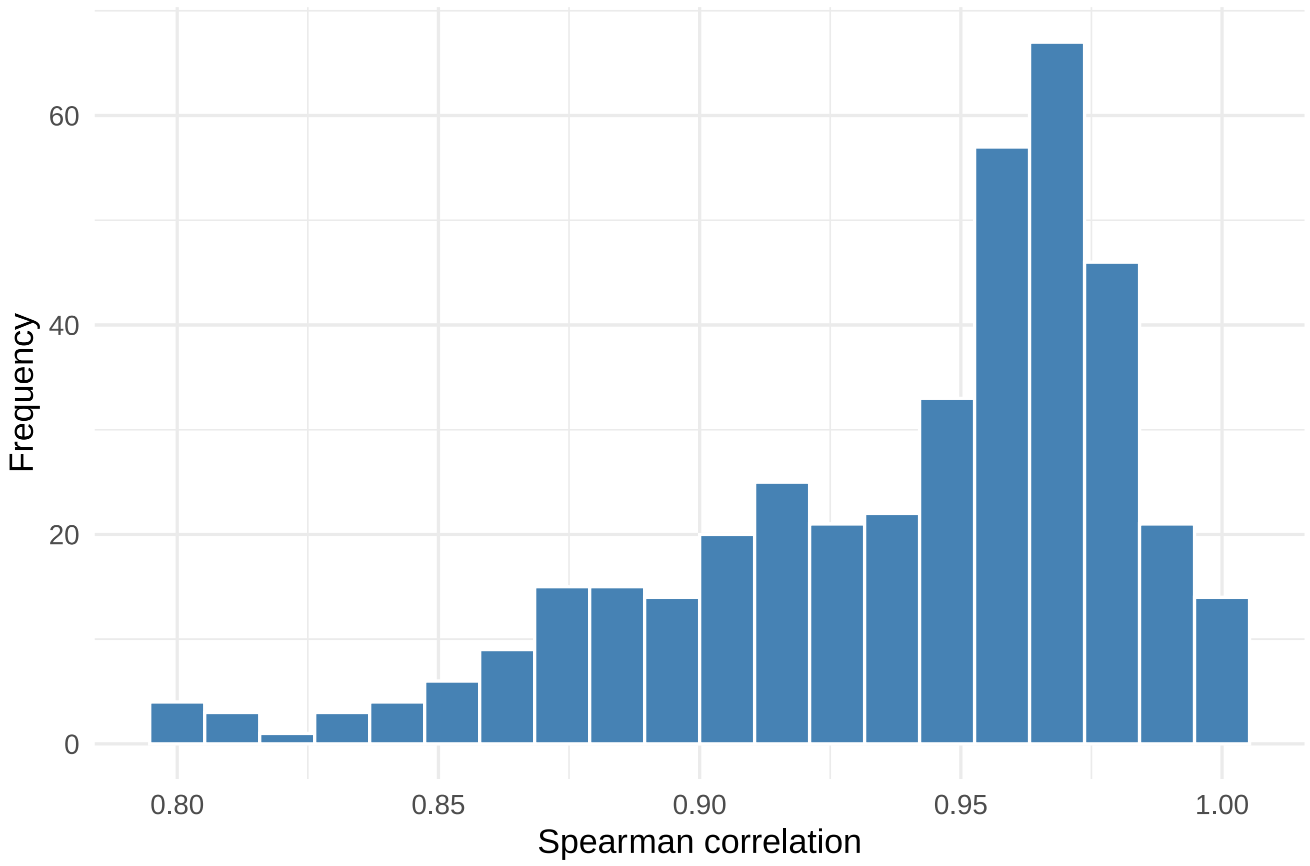


**Figure S3.** Reproducibility of LLM-based pipeline ranking. Histogram of Spearman correlations between pipeline rankings obtained from repeated runs of Lense on identical datasets. Each correlation is computed between pairs of runs.

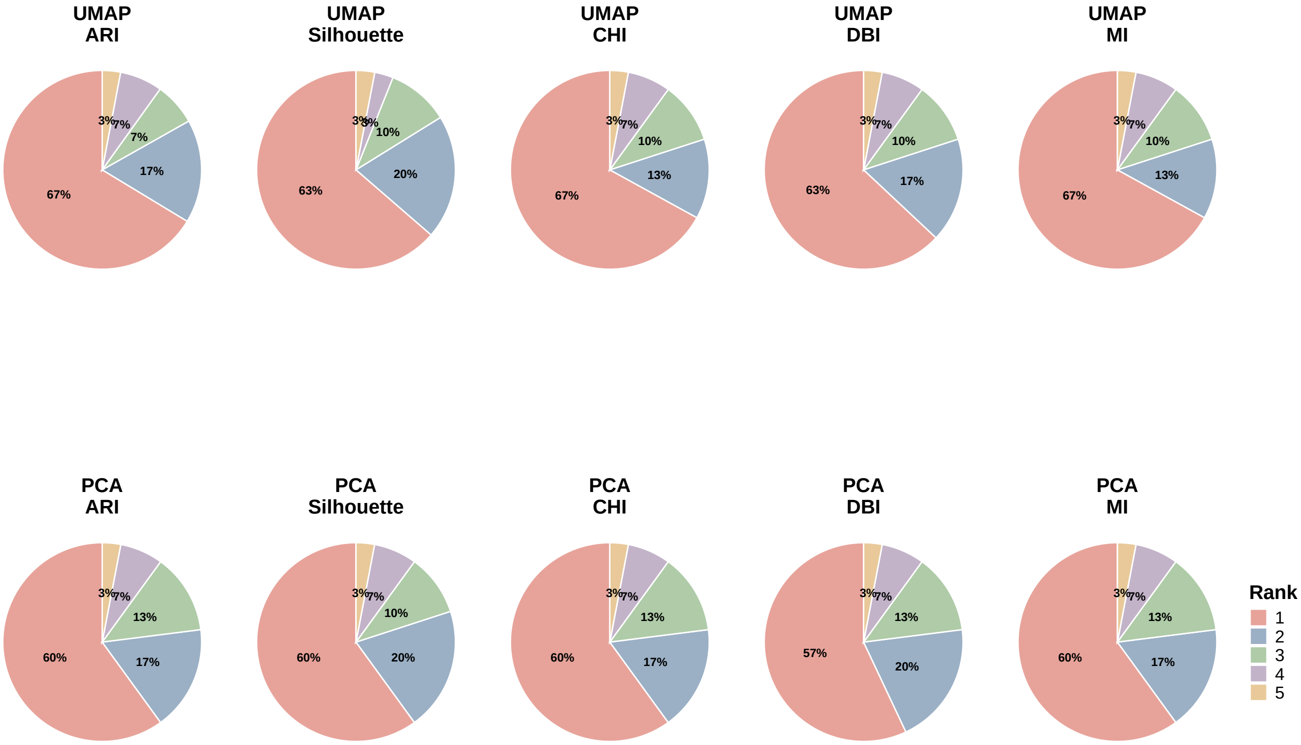


**Figure S4.** Proportion of simulation scenarios in which Claude achieved different ranks. Pie charts show the distribution of ranks selected by Claude across simulation scenarios under different evaluation metrics (ARI, Silhouette, CHI, DBI, and MI) and embedding methods (UMAP and PCA).

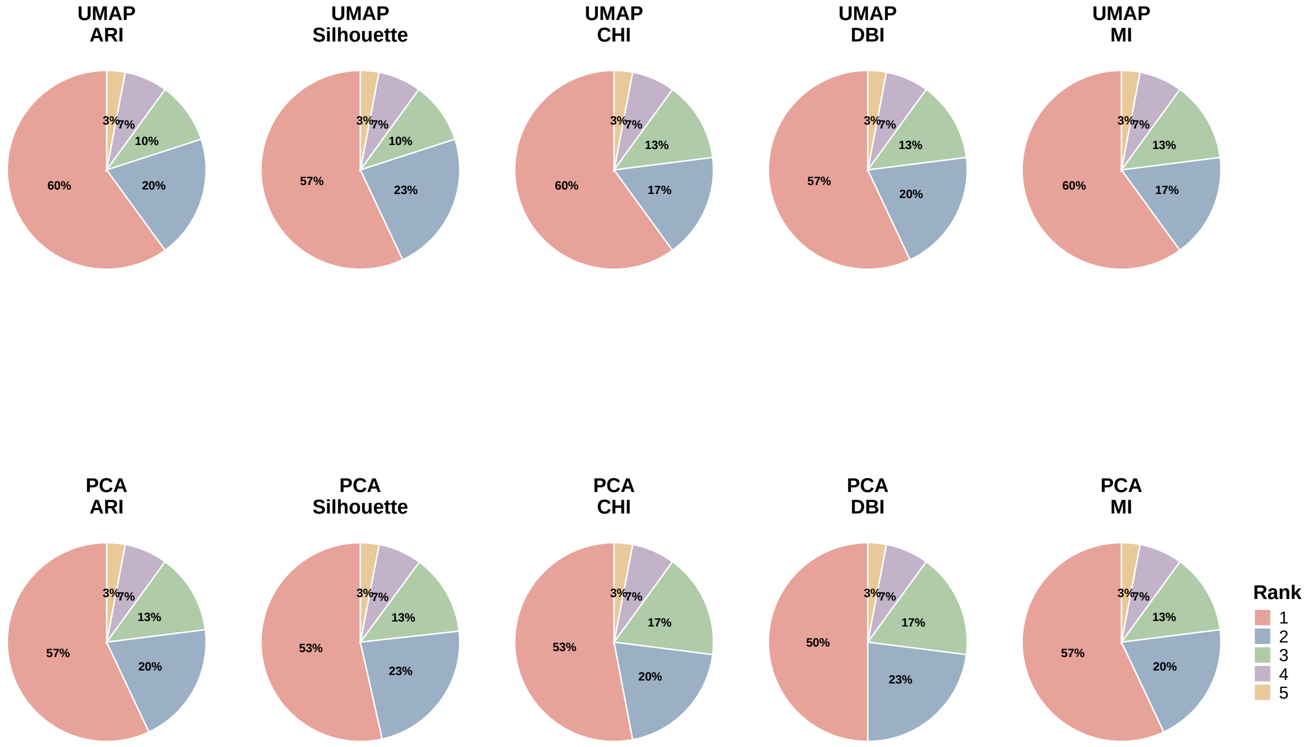


**Figure S5.** Proportion of simulation scenarios in which Gemini achieved different ranks. Pie charts show the distribution of ranks selected by Gemini across simulation scenarios under different evaluation metrics (ARI, Silhouette, CHI, DBI, and MI) and embedding methods (UMAP and PCA).

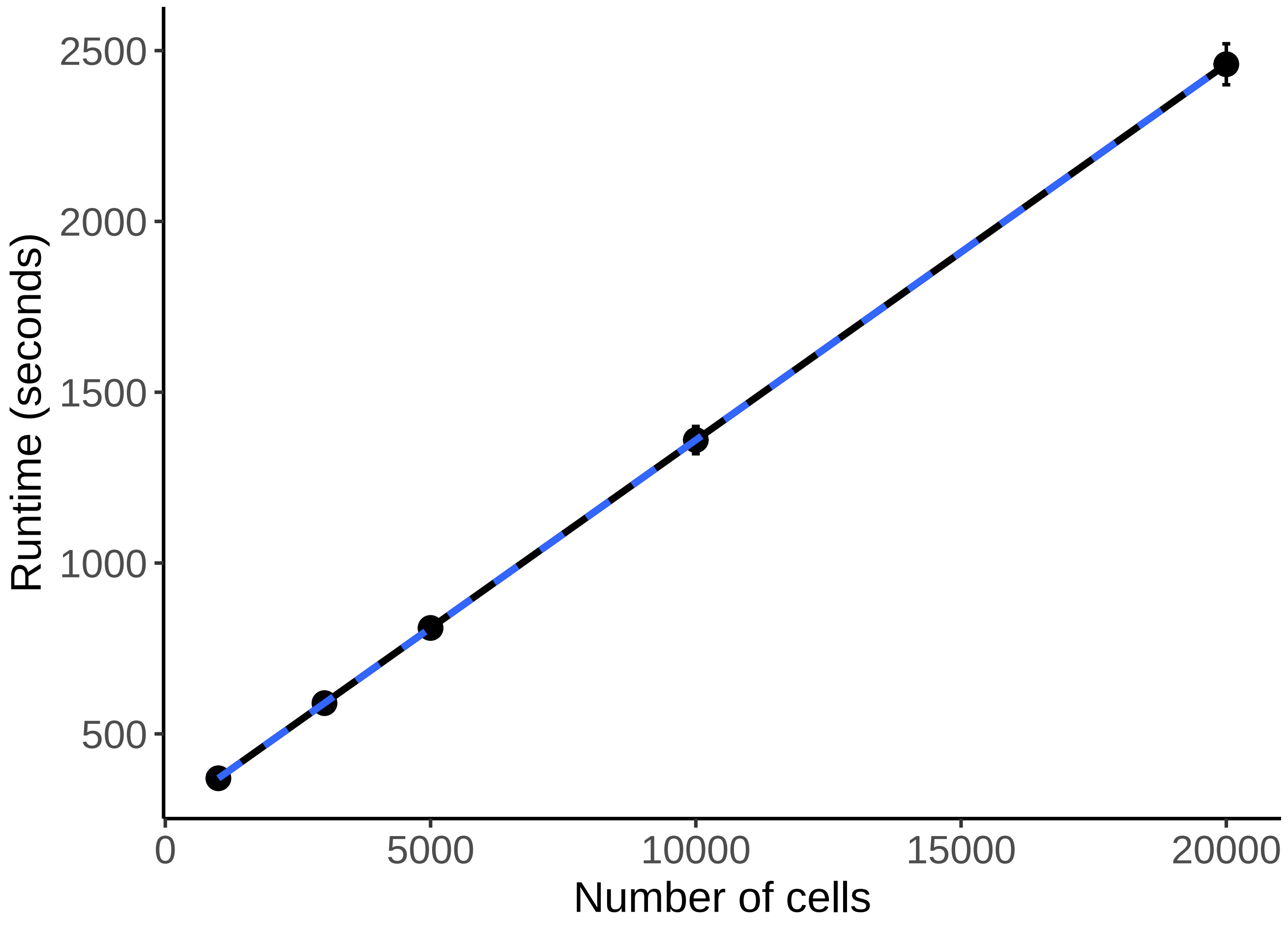


**Figure S6.** Computational performance of Lense, showing runtime (in seconds) as a function of the number of cells.
